# Subunit specialisation in AAA+ proteins and substrate unfolding during transcription complex remodelling

**DOI:** 10.1101/2025.02.15.638424

**Authors:** Forson Gao, Fuzhou Ye, Martin Buck, Xiaodong Zhang

## Abstract

Bacterial RNA polymerase (RNAP) is a multi-subunit enzyme that copies DNA into RNA in a process known as transcription. Bacteria use σ factors to recruit RNAP to promoter regions of genes that need to be transcribed, with 60% bacteria containing at least one specialized σ factor σ^54^. σ^54^ recruits RNAP to promoters of genes associated with stress responses and forms a stable closed complex that does not spontaneously isomerize to the open state where promoter DNA is melted out and competent for transcription. The σ^54^-mediated open complex formation requires specific AAA+ proteins (<u>A</u>TPases <u>A</u>ssociated with diverse cellular <u>A</u>ctivities) known as bacterial enhancer-binding proteins (bEBPs). We have now obtained structures of new intermediate states of bEBP-bound complexes during transcription initiation, which elucidate the mechanism of DNA melting driven by ATPase activity of bEBPs and suggest a mechanistic model that couples the ATP hydrolysis cycle within the bEBP hexamer with σ^54^ unfolding. Our data reveal that bEBP forms a non-planar hexamer with the hydrolysis-ready subunit located at the furthest/highest point of the spiral hexamer relative to the RNAP. ATP hydrolysis induces conformational changes in bEBP that drives a vectoral transiting of the regulatory N-terminus of σ^54^ into the bEBP hexamer central pore causing the partial unfolding of σ^54^, while forming specific bEBP contacts with promoter DNA. Furthermore, our data suggest a mechanism of AAA+ protein that is distinct from the hand-over-hand mechanism proposed for many AAA+ proteins, highlighting the versatile mechanisms utilized by the large protein family.

## Introduction

Transcription is an essential cellular process carried out by RNA polymerases (RNAP) which are recruited to genes by a range of regulatory factors (1, 2). In bacteria, σ factors are general transcription factors that recruit RNAP to specific promoter regions upstream of the transcription start site (TSS, +1) (3). Initially, the RNAP-σ holoenzyme forms a transcription-incompetent closed complex (RPc) with the duplexed promoter DNA (dsDNA), which is not accessible to the RNAP catalytic site. Transcription initiation involves converting the dsDNA into single-stranded DNA and loading the single template strand to the active site to form the transcription-competent open complex (RPo) (4, 5). In bacteria, most transcription is initiated by the housekeeping σ^70^ factor, which recruits RNAP to the -35 and -10 consensus regions of promoters (upstream relative to TSS) (6). In most cases σ^70^ can spontaneously nucleate DNA melting as well as stabilize, propagate and load the DNA transcription bubble into the RNAP active site without the need for transcription activators (7).

σ^54^ is a major variant σ factor that recruits RNAP to genes associated with nutrient depletion, infection and other stress response (3). σ^54^ is structurally and mechanistically distinct from σ^70^ (8). σ^54^ forms a very stable closed complex upon binding to distinct -24(GG) and -12(GC) regions of the promoter, and requires a specialized class of transcription activator proteins that bind DNA 100-150 bp upstream from TSS, thus called bacterial enhancer-binding proteins (bEBPs) (9). bEBPs belong to the large <u>A</u>TPase <u>A</u>ssociated with diverse cellular <u>A</u>ctivities (AAA+) family of proteins and function as hexamers (10). DNA looping enables bEBP hexamers to engage and actively remodel the RNAP-σ^54^-DNA closed complex to convert it into an open complex (4).

Our previous work has revealed how σ^54^ forms a stable closed promoter complex, preventing DNA from entering the RNAP and therefore tightly inhibiting transcription (11, 12). bEBP binding induces conformational changes in σ^54^ and RNAP, eventually leading to open complex formation (12). Our recent work has provided mechanistic understanding on how bEBP interacts with σ^54^ and DNA (13). Strikingly, we observed that the σ^54^ uses its N-terminal helix (RI-H1) to “pry” open the promoter DNA, allowing the N-terminus of σ^54^ to thread through the opening in the promoter DNA, before being captured and enclosed by bEBP hexamer in its central pore (**Fig. 1a**) (13). Furthermore, there are extensive interactions between the bEBP and promoter DNA. However, how ATP binding and hydrolysis amongst the six subunits of an bEBP promote conformational changes in σ^54^ and DNA that lead to the relief of σ^54^ inhibition and full transcription bubble formation remains unknown (13).

**Fig. 1.**
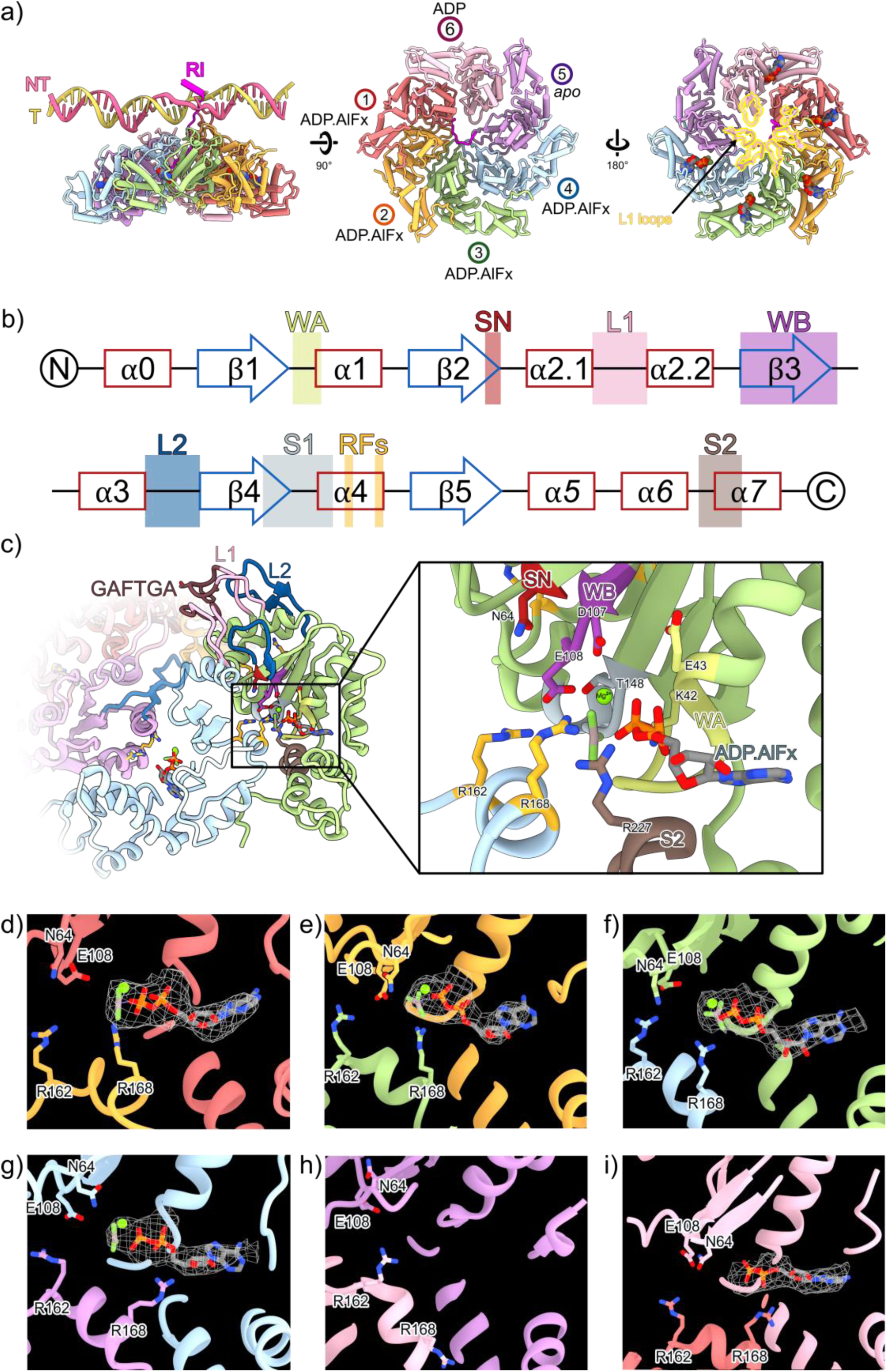
Structural analysis of bEBP hexamer in the transcription intermediate state of RNAP-σ^54^-PspF_1-275_ with DNA mismatched at -12/-11. (a) overview of the hexamer and its arrangement with DNA and σ^54^ RI. (b) functional motifs and secondary structures of bEBP AAA+ domains. (c) nucleotide binding pocket, with key residues in nucleotide binding and hydrolysis labelled. (d)-(i) density corresponding to nucleotides in each protomer with key catalytic residues labelled.

Previously we have identified that the signature sequence motif of bEBPs (GAFTGA) is within a surface loop inserted into the AAA+ ATPase domain (14). Our recent work shows that this loop is responsible for enclosing the σ^54^ N-terminal peptide (13). High resolution crystal structures of the bEBPs phage shock protein F (PspF) (15) and those of nitrogen regulator protein C (NtrC1) in complex with different nucleotides (16) showed that ATP-bound form correlated with L1 loop in an extended conformation while ADP-bound form associated with a folded down conformation (15). Based on these studies and those of other AAA+ proteins, it was proposed that a particular nucleotide state controls L1 loop conformations while L1 loop conformation is communicated back to the nucleotide-binding site via a signalling network, with an extended L1 loop allowing ATP hydrolysis (17). Our recent work suggests that nucleotide hydrolysis is coupled to σ^54^ N-terminal translocation via bEBP L1 loops.

In this work, we analyse our previously resolved cryo electron microscopy (cryoEM) reconstruction of PspF-RNAP-σ^54^ in complex with promoter DNA with mismatched bases at -12/-11 (RPi(-12/-11)), mimicking the DNA fork junction structure seen in RPc, to reveal the precise functional states of PspF subunits within the hexamer ring. Correlating these functional states with σ^54^ interactions, we propose a mechanistic model that couples the ATP hydrolysis cycle within the bEBP hexamer with σ^54^ N-terminal translocation. Using promoters that are partially melted (between -11 and -8) and fully melted (between -10 and -1) to mimic a partially or fully established transcription bubble, we determined the structures of bEBP-bound transcription complexes using cryoEM. These structures allowed us to capture additional intermediate states which provide further insights into how DNA melting is coupled to bEBP hexamer conformation and how ATP hydrolysis drives σ^54^ unfolding.

## Results

### Analysis of PspF_1-275_ hexamer in RPi(-12/-11) suggests a molecular mechanism for ATP hydrolysis coordination within the hexamer

Detailed inspection and analysis of RPi reconstruction revealed clear electron density to allow us to unambiguously determine the nucleotide states of each protomer (**Fig. 1**). Based on the nucleotide density, we assigned protomers 1-4 to have Mg^2+^-ADP.AlFx bound (**Fig. 1a, d-g**), whereas protomer 5 empty (*apo*) and protomer 6 ADP-bound (**Fig. 1h-i**). Inspecting each nucleotide binding pocket, we could identify key residues that are involved in nucleotide binding and hydrolysis including the Walker A, Walker B and sensor 2 motifs *in cis*, with *in trans* interactions from R-fingers (R162, R168) present in the adjacent protomer (**Fig. S1a-f**). Previous studies on AAA+ ATPases have revealed that R-fingers, Walker B (D107 and E108) and the associated “glutamate-switch” residue (N64) are required to be positioned in correct conformation to enable ATP hydrolysis (ATP hydrolysis-ready state) (17, 18). Comparing positions of these catalytic residues suggest that protomers 1 to 4 have *trans* R-fingers correctly positioned for ATP binding (**Fig. S1**). Indeed protomers 1-4 have similar conformations (**Fig. S2a-b**). Interestingly only protomer 1 has clear density showing the conformation of Walker B (E108) and the corresponding “glutamate-switch” (N64) in hydrolysis-ready conformation (17) (**Fig. S1**). In protomers 5 and 6, the α-lid is further closed relative to αβ sandwich subdomain, correlating with their nucleotide states of apo and ADP respectively (**Fig. S2c**).

There are extensive interactions from protomer 1 through to protomer 4 (**Figs. 1a, 2a**). Indeed, L1 and L2 in protomers 2-4 are constrained, adopting the folded down conformation (**Fig. 2, Fig. S2b**). The conserved phenylalanine on the GAFTGA loop of protomer 1 sterically blocks the L1 loop of protomer 2 from raising upwards (**Fig. 2b**). The interfaces between protomers 1 and 6, between protomers 6 and 5, are smaller compared to other dimer interfaces in the hexamer (**Fig. 1a**), suggesting that protomers 1, 5 and 6 are less constrained inside the hexamer. Viewed from the side, PspF protomers 1-5 form a spiral with protomer 1 at the highest point (closest to σ^54^ RI) while protomer 5 at the lowest point. Protomer 6 is between protomer 1 and protomer 5 (**Figs. 1a, 2a**).

**Fig. 2.**
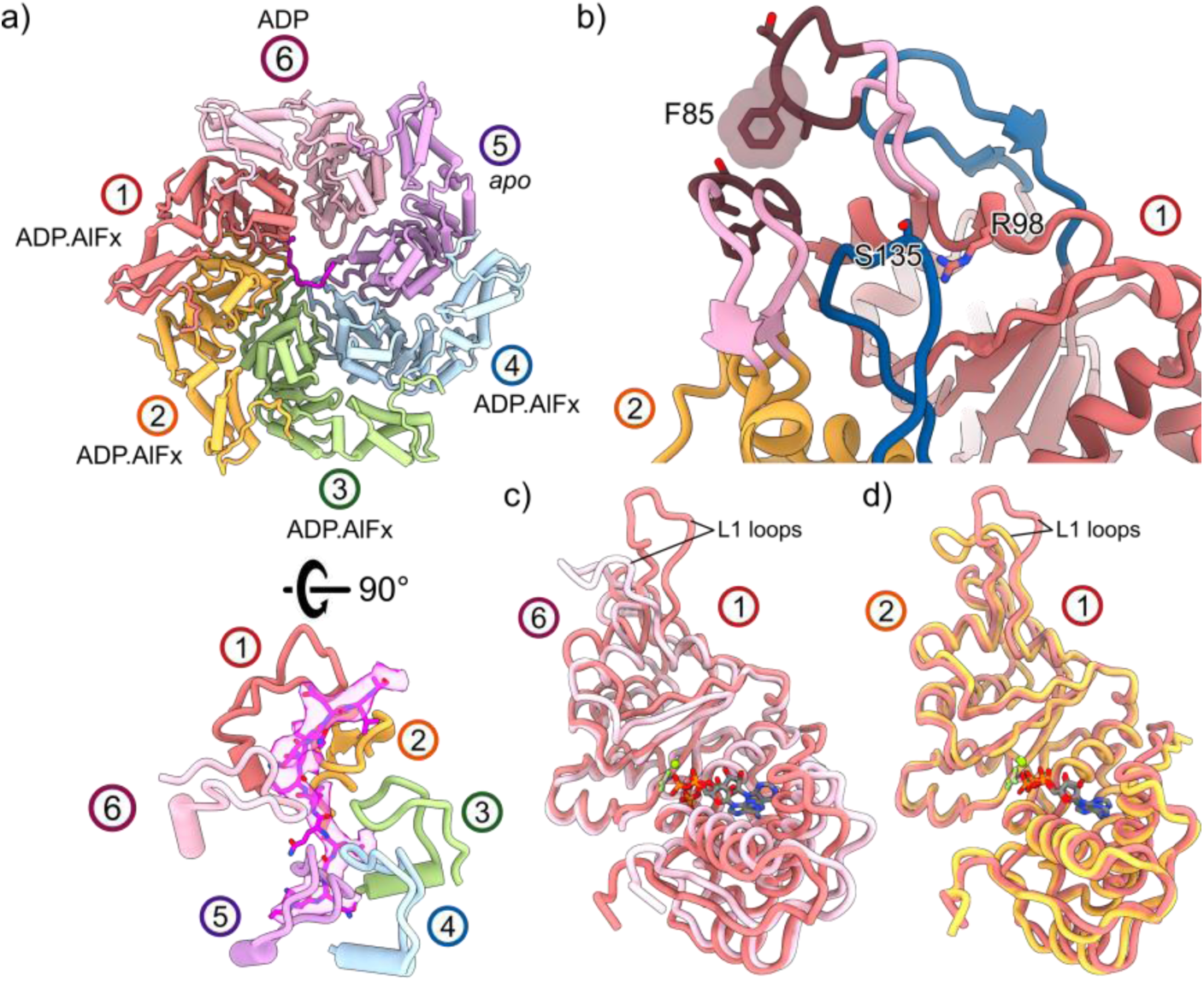
Protomer conformations and their functional state. (a) L1 loops of protomers track σ^54^ N-terminus in the hexamer pore with protomer 1 at the higher position in the helical spiral. (b) conformations and interactions of L1 (brown) and L2 (blue) loops between protomer 1 and protomer 2. (c) Overlays between protomer 1 (ATP*) and protomer 6 (ADP) reveal the conformational changes upon ATP hydrolysis. (d) overlay of protomer 1 and protomer 2 shows the overall similarities in their conformation and different conformations of the L1 loops.

The configuration of nucleotide binding pockets of the PspF_1-275_ hexamer suggests that protomer 1 is likely the subunit to hydrolyse ATP next, becoming an ADP-bound state. Comparing the structures of protomer 1 (ATP state) and protomer 6 (ADP state) reveals the conformational changes in PspF_1-275_ protomer upon hydrolysis (**Fig. 2c**). In addition to the subdomain closure when transitioning from protomers 1 to 6, we also observe that the L1 loop folds downward from an extended conformation (**Fig. 2c**, **Fig. 3a**).

**Fig. 3.**
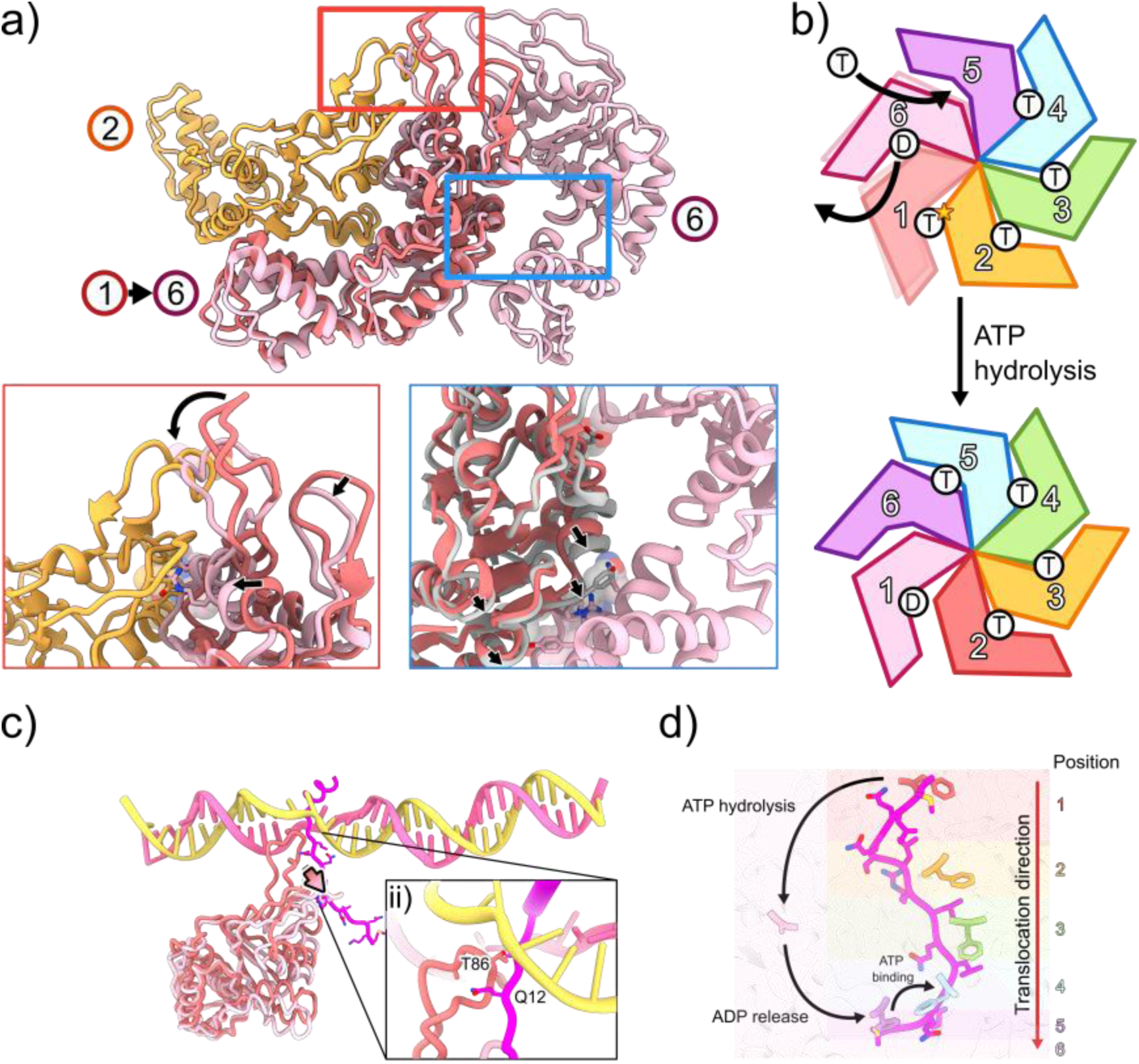
Proposed ATP hydrolysis cycle and conformational changes within PspF_1-275_ hexamer. (a) protomers 1, 2 and 6 in the hexamer and conformational changes in protomer 1 upon hydrolysis. Insets: left - ATP hydrolysis of protomer 1 would cause it to adopt conformation of protomer 6 (arrow), causing a steric clash with protomer 2; right -upon hydrolysis, protomer 1 could move (arrows) towards protomer 6. (b) proposed ATP hydrolysis and nucleotide-state changes within the hexamer. (c) ATP hydrolysis of protomer 1 would pull σ^54^ N-terminus, (d) proposed positional changes in the hexamer spiral upon ATP hydrolysis of protomer 1.

Upon hydrolysis of protomer 1, the conformational changes in protomer 1 would cause steric clashes with protomer 2 at multiple positions on the ⍺β sandwich subdomain, as well as the GAFTGA motif within the L1 loop (**Fig. 3a, insets, arrows**). Protomer 2 is tightly packed against protomer 3 while the interface between protomer 1 and protomer 6 is looser (**Fig. 3a**). This suggests that upon hydrolysis and to avoid steric clashes, protomer 1 is likely to detach from protomer 2 and so moves towards protomer 6 (**Fig. 3a)**. The detachment of protomer 1 from protomer 2 would relieve the constraint on L1 loop in protomer 2 (**Fig. 2b**), allowing the L1 loop to be extended, reaching to interact with σ^54^ RI, adapting new position previously occupied by protomer 1 (**Fig. 3b**). The extended L1 loop would also reposition the nucleotide binding pocket, making it hydrolysis ready (15, 17, 19).

We propose that the detachment of protomer 1 from 2 not only enables protomer 2 to adapt hydrolysis-ready and σ^54^ RI engaged state, protomer 1 would now engage with protomer 6, promoting nucleotide release of protomer 6 (becoming apo state). Presumably this then causes conformational changes in protomer 6 and subsequently protomer 5, which in turn can now bind ATP, completing the shift of nucleotide states within the hexamer by one subunit (**Fig. 3b**).

The proposed mechanism suggests the power stroke occurs when protomer 1 hydrolyses ATP to adapt the position occupied by protomer 6. L1 loop in protomer 1 has the most extensive interactions with σ^54^ Region I and promoter DNA. Upon hydrolysis, the L1 loop folds downwards, and this movement would thus pull σ^54^ N-terminal peptide with it (**Fig. 3c**). The remaining PspF_1-275_ protomers form primarily hydrophobic “slippery” interactions with conserved hydrophobic residues on σ^54^ N-terminal peptide and could enable the sliding of the rest of the peptide (**Fig. 3d**). The translocation of N-terminus would cause unfolding of RI-H1, which has been shown to be crucial in maintaining the stable closed promoter complex (12, 13).

Significantly, L1 loops make extensive interactions with promoter DNA (**Fig. 3c**), thus the changes in L1 loop conformation within the hexamer would change interactions with DNA. Further, the RI translocation would also change its interactions with DNA. These altered interactions likely contribute to the DNA melting induced by PspF ATP hydrolysis.

### Structures of RPi(-11/-8) and RPi(-10/-1) complex

To test this model and to further probe how ATP hydrolysis induces or stabilizes changes in promoter DNA, we used synthetic DNA with differing degrees of pre-melted DNA to mimic melting intermediates during transcription bubble formation (**Fig. S3a**). Stable complexes can be detected with all the DNA substrates tested (**Fig. S3b**). Some of the substrates could support transcription with dATP or ADP.AlFx (**Fig. S3c**), suggesting that stable intermediate complexes formed on these substrates can proceed to open complex formation. Interestingly, DNA with a partially opened bubble between -11 and -8 did not support transcription (**Fig. S3c**), suggesting that this complex is unable to proceed further to open complex. To understand the structural basis and how DNA is stabilised in these complexes, we determined the structures of PspF_1-275_-RNAP-σ^54^-DNA containing mismatches between -11 and -8 (RPi(-11/-8)) (**Figs. S4-6, Table S1**) and between -10 and - 1 (RPi(-10/-1)) (**Figs. S7-8, Table S1**). These structures reveal new transcription complex conformations not previously observed.

In the RPi(-11/-8), we observe multiple structural conformations, at varying degrees of tilt of PspF_1-275_ hexamer ring relative to RNAP-σ^54^ and varying degrees of RNAP clamp opening (**Figs. S4, S6**) We also observe significant degrees of promoter DNA bending, demonstrating the wide degree of conformational flexibility in the complex (**Fig. S4**). The DNA density downstream of -10 was generally poorly resolved, indicating that this region was highly flexible. However, one conformation has sufficiently high resolution to allow us to resolve the DNA (**Fig. S9b**). In this conformation, we observe the PspF hexamer ring is tilted by ∼50° compared to that of RPi(-12/-11) and the DNA is bent by ∼45° (**Fig. 4**)

**Fig. 4.**
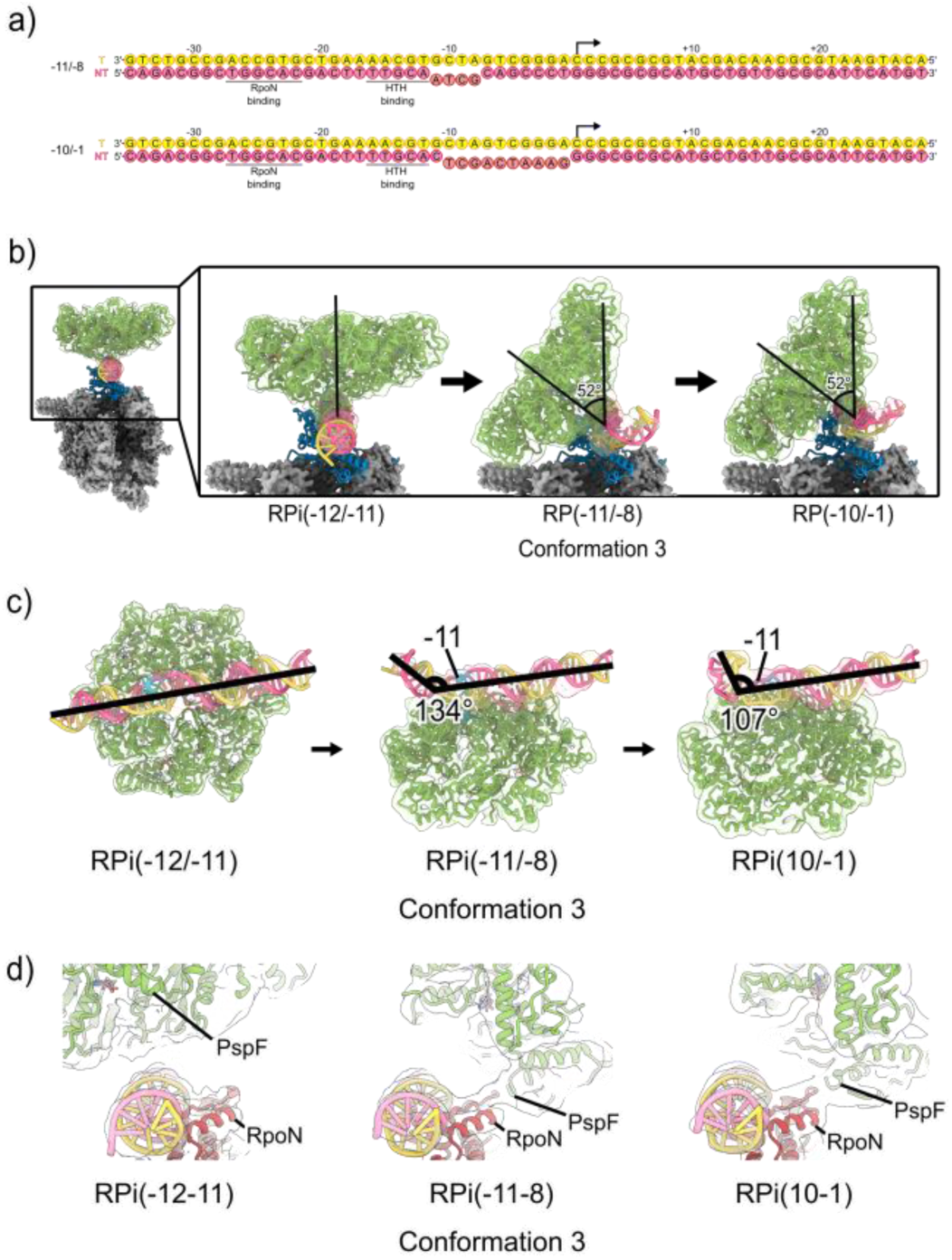
Structures of RPi (-11/-8) and RPi (-10/-1). (a) DNA substrates used to mimic partially or fully melted transcription bubble. (b) comparisons of the PspF_1-275_ hexamer in relation to RNAP in the RPi complexes. (c) comparisons of DNA trajectories in these complexes. (d) additional interactions between PspF_1-275_ and σ^54^ RpoN domain are observed in RPi(-11/-8) and RPi(-10/-1).

In the RPi(-10/-1) complex, we also observe a range of PspF hexamer ring conformations with one resolved to higher resolution (**Fig. S7**). Interestingly, in this conformation, the hexamer ring has similar tilt as that of RPi (-11/-8) at ∼50° while the DNA is bent even further (>70°) compared to that RPi(-11/-8) (**Fig. 4**).

Despite the conformational differences, similarly to RPi(-12/-11) structure, the protomer that engages with DNA is also at the highest position within the hexamer ring (**Fig. 5a-b**). In these complexes, we observe a previously unreported interaction between PspF and σ^54^ RpoN domain (**Fig. 4d**). This interaction is only possible with the tilted ring conformation, suggesting this interaction contributes to the stabilisation of this ring conformation in both RPi(-11/-8) and RPi(-10/-1), giving rise to the higher resolution reconstruction of these conformations.

**Fig. 5.**
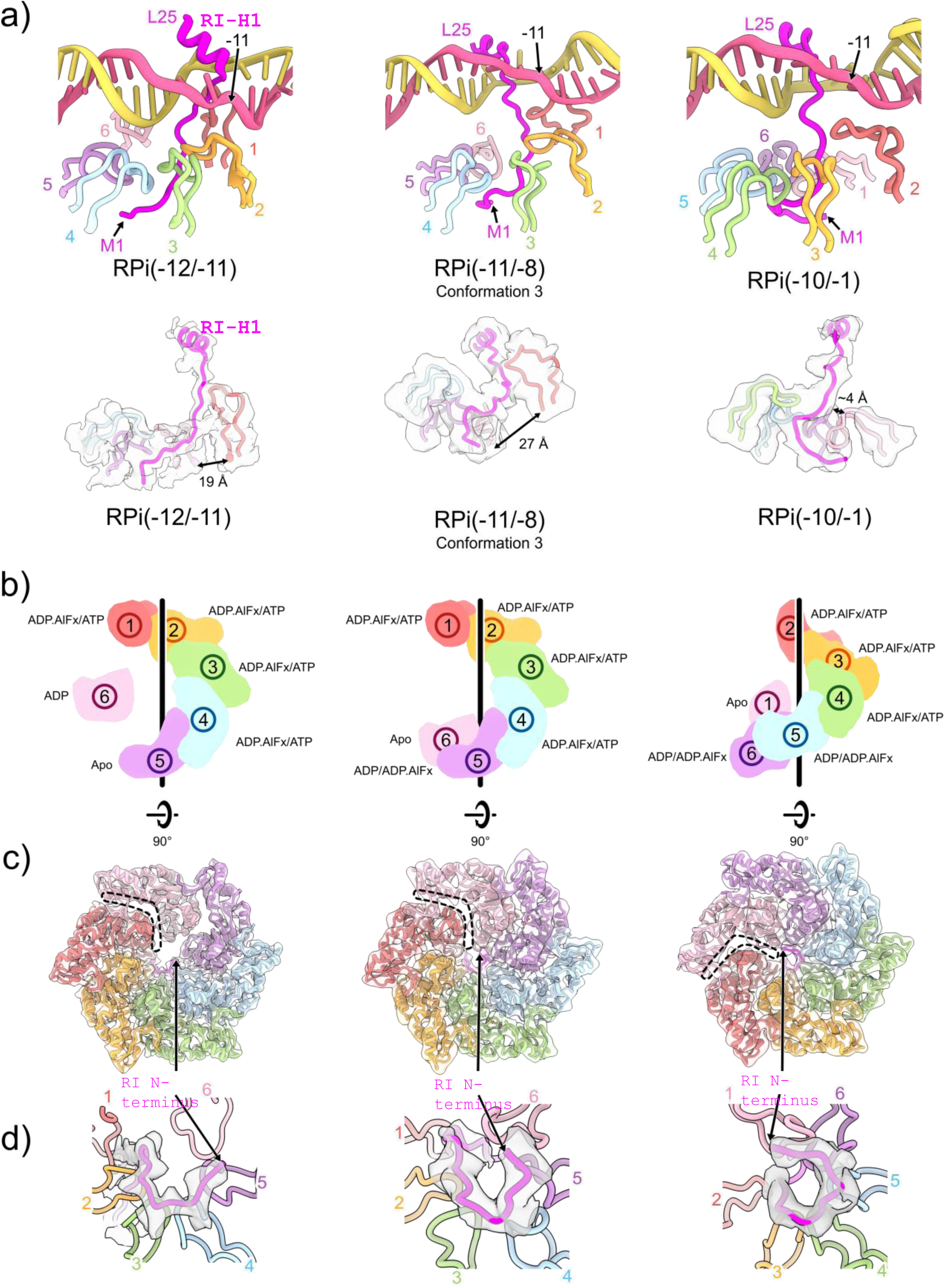
ATP hydrolysis couples to σ^54^ N-terminus translocation and RI-H1 unfolding. (a) PspF_1-275_ hexamer is arranged differently in these complexes, showing as the distances between equivalent elements in protomer 1 and protomer 6. Density for RI-H1 is also progressively worse from RPi(-12/-11) to RPi(-11/-8) and RPi(-10/-1), (b) cartoon depictions of the protomers in the hexamer spiral. (c) top views of the hexamer showing the relative positions of the protomers in the spiral. Dashed lines indicate boundaries between protomers 1 and 6. (d) density for σ^54^ N-terminus, indicating that N-terminus has been translocated further into the hexamer from RPi(-12/-11) to RPi (-11/-8) and RPi (- 12/-11).

Given the PspF_1-275_ rings are in similar positions between RPi(-11/-8) and RPi(-10/-1) complexes, we could compare the relative positions of each protomer between RPi(-11/-8) and RPi(-10/-1) (**Fig. 5c**). Compared to RPi(-11/-8), the position of the power stroke protomer (protomer 1) has shifted within the ring, with the protomer in position 2 now becomes the power stroke protomer (protomer 1 position) in (**Fig. 5b-c**), suggesting that these two complexes could represent two states as described in **Fig. 3** after the hydrolysis of protomer 1. Interestingly, protomer 1 in RPi(-10/-1) is shifted significantly towards protomer 6 compared to that of RPi(-11/-8), with the distance between L1 of these two protomers reduced from 27 Å to 4 Å (**Fig. 5a**), consistent with our proposed mechanistic model that hydrolysis of protomer 1 at the top of the spiral causes it to relocate downwards in the hexamer spiral (**Fig. 5b**). We could not overlay the hexamers between RPi(-12/-11) with RPi(-11/-8) due to the significant tilt and rotation of the hexamer. However, based on the position of the highest protomer and its interactions with DNA, we could assign relative numbering within the hexamer. Comparing the relative positions suggest that protomers 1-5 are similar while protomer 6 is shifted towards protomer 5 (**Fig. 5b**). Together with RPi(-12/-11), we capture three conformational states (**Fig. 5b**) that are consistent with the model proposed in **Fig. 3d** in terms of protomer repositioning during ATP hydrolysis cycle.

Looking at the density within the PspF hexamer, we observe density that corresponds to σ^54^ N-terminus (**Fig 5d**). Significantly, we observe the σ^54^ N-terminus is further shifted along the PspF hexamer in RPi(-11/-8) and RPi(-10/-1) compared to that of RPi(-12/-11) (**Fig. 5c**). This would thus support the mechanistic model in which we proposed that ATP hydrolysis and the conformational changes within the hexamer lead to translocation of the σ^54^ N-terminus; and RPi(-11/-8) and RPi(-10/-1) are further along the transcription initiation pathway. Indeed, we observe reduced quality density for RI-H1 (**Fig. 5a**), consistent with RI N-terminal translocation leading to the unfolding of RI-H1, eventually releasing the constraint on σ^54^ ELH.

## Discussion

Our results strongly support a molecular mechanism by which bEBPs utilise ATP hydrolysis to thread σ^54^ N-terminus through the nascent transcription bubble into the hexamer pore of AAA+ domains. Upon ATP hydrolysis, there are two consequences: firstly, σ^54^ N-terminus is further translocated into the AAA+ hexamer pore. Translocation of σ^54^ N-terminus leads to the unfolding of RI-H1 helix, disrupting the hydrophobic interactions between RI-H1 and ELH. Given that RI-H1-ELH plays crucial roles in suppressing transcription initiation, unfolding R1-H1 (∼ 26 residues) instead of the whole peptide chain should be sufficient to relocate ELH and relieve the inhibition. Secondly, PspF_1-275_ hexamer ring tilts, as observed in RPi(-11/-8) and RPi(-10/-1) structures, altering interactions between PspF_1-275_ and DNA, which could contribute to or stabilise DNA melting. Further, PspF ring tilting directly influences DNA conformation as the observed DNA bending in RPi(-11/-8) and RPi(-10/-1). DNA bending would further promote DNA melting and loading into the RNAP active site (**Fig. 4**).

Our structures of PspF_1-275_ in the RPi complexes suggest that ATP binding occurs in the protomer that is located at the bottom of a hexamer spiral and protomers are increasingly positioned to be hydrolysis-ready anti-clockwise, with protomer 1 at the top of the spiral performing the power stroke by hydrolysing ATP while translocating the N-terminus of σ^54^, before transitioning to conformation of protomer 6, while protomer 2 becomes the next power stroke subunit (**Fig. 3b**, **Fig. 5b**). This is in contrast with the sequential model proposed for several AAA+ proteins. These AAA+ proteins use their pore loops to engage with their substrates (20). Structural studies including RuvB (21), Hsp104 (22), Lon protease (23), p97, Vps4 (24) and FIGNL1 (24) have revealed that conserved aromatic residues of the pore loops, arranged in a spiral staircase, engage with substrates. In PspF, L1 loops instead of pore loops are utilized in substrate engagement. The equivalent pore loops in PspF do not have the conserved KW/Y motifs at the tip and are located too far from the substrate (**Fig. S10**). In this model, ATP hydrolysis in protomer 4 drives the translocation of substrate and promotes nucleotide release in protomer 5, results protomer 5 to detach from the substrate. Upon nucleotide exchange, the protomer re-engages, and occupies the position at the top of the spiral (20).

A further model has been proposed for ClpX. In this model, the power stroke is performed by ATP hydrolysis of protomer 1 at the top of the spiral, which shifts its position in the hexamer from position 1 towards the bottom of the spiral, and so translocating the substrates downwards. Our proposed model here for bEBPs is more consistent with this model (25). Further unlike bEBPs do not translocate peptides continuously to unfold a protein/domain, instead it partially unfolds a protein/domain. It is unknown how many nucleotide hydrolysis will be required or how the hydrolysis and unfolding are regulated.

## Acknowledgements and funding sources

Initial screening of electron microscopy grids was carried out at Imperial College London Centre for Structural Biology. We acknowledge Diamond Light Source for access and support of the cryoEM facilities at the UK national eBIC, proposal EM19865, funded by the Wellcome Trust and MRC, and London Consortium for high resolution cryoEM (LonCEM), funded by the Wellcome Trust. This project is funded by the UKRI to X.Z. and M.B. (BB/M011178/1). F.G. is funded by a BBSRC DTP studentship.

## Materials and methods

### Protein purification

All chromatography was performed on a ÄKTA Pure (Cytiva^TM^) at 4 °C. Flow rates and method details for each chromatography run can be found in Supplementary Table 1). After each chromatography run, fractions from selected peaks were then ran on Bolt 4-12% Bis-Tris Plus SDS-PAGE gels alongside PageRuler Plus Prestained 10 to 250 kDa molecular weight ladder (Thermofisher Scientific™). All SDS-PAGE gels were run at 200 V for 35 minutes before being stained with InstantBlue (Expedeon^TM^) and imaged with Gel Doc™ XR+ Gel Documentation System (BioRad^TM^).

#### σ^54^ purification

σ^54^ was initially extracted from lysate by nickel affinity chromatography, with a 5 ml HisTrap™ HP column (Cytiva^TM^) equilibrated with σ^54^ NiA buffer (20 mM Tris-HCl pH 8, 500 mM NaCl, 5% v/v Glycerol, 10 mM Imidazole) and eluted with a 5-100% linear gradient of σ^54^ NiB buffer (70 mM NaCl, 500 mM imidazole pH 8, 5% v/v glycerol). Following SDS-PAGE analysis, selected fractions were pooled and passed through a HiPrep™ 26/10 desalting column (Cytiva^TM^) to exchange into HepA buffer (20 mM Tris-HCl pH 8, 50 mM NaCl, 5% v/v glycerol, 1 mM TCEP).

σ^54^ was then further purified by heparin affinity chromatography with a 5 ml HiTrap™ Heparin HP column (Cytiva^TM^) equilibrated with HepA buffer and eluted with a 5-100% gradient of HepB buffer (20 mM Tris-HCl pH 8, 1 M NaCl, 5% v/v glycerol, 1 mM TCEP). Fractions were pooled based on SDS-PAGE analysis.

Pooled fractions were then gel filtrated using a Superdex Hiload® 75 16/600 column (Cytiva^TM^) equilibrated with GF buffer (20 mM Tris-HCl pH 8, 150 mM NaCl, 5% v/v glycerol, 2 mM TCEP). Fractions were pooled based on SDS-PAGE analysis and concentrated using a 30 kDa molecular weight cut-off (MWCO) centrifugal filter (Merck Millipore). The concentration was measured using a Nanodrop ND-1000 spectrophotometer (Thermofisher Scientific™). Protein was then aliquoted into 20 μl volumes, flash frozen in liquid nitrogen and stored at -80 °C.

#### PspF_1-275_ purification

PspF_1-275_ was initially extracted from lysate by nickel affinity chromatography; clarified lysate was passed through a 5 ml HisTrap HP column (Cytiva^TM^) equilibrated with PspF NiA buffer and eluted with 5-100% PspF NiB buffer (50mM NaH_2_PO_4_, 50 mM NaCl, 500 mM imidazole pH 8, 5% glycerol). The peak fractions were pooled, concentrated to less than 5 ml using a centrifugal filter and dialysed in PBS for 1 hour at room temperature with 10 kDa MWCO SnakeSkin™ dialysis tubing (Thermofisher Scientific™).

The N-terminal 6×His tag was then cleaved by adding 90 μl (18 NIH units) thrombin. The dialysis tubing was transferred to a fresh beaker of PBS for 2 hours at room temperature. Complete cleavage was verified by taking periodic aliquots for SDS-PAGE analysis and running for 50 minutes. Any uncleaved PspF_1-275_ was then removed by reverse nickel affinity chromatography, in which the flowthrough was collected for gel filtration.

Flow through fractions were gel filtrated using a Superdex Hiload® 75 16/600 column (Cytiva^TM^) equilibrated with GF_PspF_ buffer (10 mM Tris-HCl pH 8, 50 mM NaCl, 5% v/v glycerol, 0.1 mM EDTA, 1 mM TCEP). Fractions were pooled based on SDS-PAGE analysis and concentrated using a 10 kDa molecular weight cut-off (MWCO) centrifugal filter (Merck Millipore^TM^). The concentration was measured using a Nanodrop ND-1000 spectrophotometer (Thermofisher Scientific™). Protein was then aliquoted into 20 μl volumes, flash frozen in liquid nitrogen and stored at -80 °C.

#### RNAP core (α_2_ββ’ω)

RNA polymerase was initially extracted from cell lysate using polymin P precipitation. Polymin P is a positively charged polymer that co-precipitates DNA-binding proteins. 10% (v/v) Polymin P was added dropwise to cell lysate at 4 °C on a magnetic stirrer to a concentration of 0.7% (v/v). The solution was then left at 4 °C for 15 minutes to homogenise, followed by ultracentrifugation at 18,000 ×*g* at 4 °C for 15 minutes. Cell pellet was then washed with 50 ml TGET buffer (10 mM Tris-HCl pH 8, 5% v/v glycerol, 1 mM EDTA, 2 mM TCEP). Centrifugation and washing were repeated twice to remove contaminants before a final wash with TGET containing 1 M NaCl to elute RNAP.

RNAP was precipitated from the eluant by adding 60% ammonium sulfate. RNAP precipitant was pelleted by ultracentrifugation as described previously and resuspended in 30 ml TGET buffer containing 100 mM NaCl. The resuspended solution was then desalted using a HiPrep™ 26/10 desalting column (Cytiva^TM^) equilibrated with 100 mM NaCl TGET buffer. Selected fractions and aliquots were collected at each stage and run on SDS-PAGE to confirm the presence of RNAP.

To further improve the purity of the RNAP, pooled fractions were then run on a BioRex 70 exchange column (BioRad^TM^), equilibrated with 100 mM NaCl TGET buffer. RNAP was eluted with an increasing gradient of 1M NaCl TGET buffer. Selected peak fractions were run on an SDS-PAGE gel before being pooled and dialysed overnight in 100 mM NaCl TGET buffer.

Pure RNAP was then obtained by HiTrap Q anion exchange chromatography, equilibrated with 100 mM NaCl TGET buffer. RNAP was eluted with a increasing gradient of 1M NaCl TGET buffer. Selected peak fractions were ran on an SDS-PAGE gel before being pooled and concentrated using a 100 kDa MWCO centrifugal filter (Merck Millipore). The concentration was measured using a Nanodrop ND-1000 spectrophotometer (Thermofisher Scientific™). Protein was then aliquoted into 20 μl volumes, flash frozen in liquid nitrogen and stored at -80 °C.

#### RNAP-σ^54^ formation

30 μM RNAP was incubated with 120 μM σ^54^ at 4 °C for 1 hour, at a final volume of 250 μl. The mixture was then loaded on a Superose 6 10/300 Increase column (Cytiva^TM^) equilibrated with GF buffer (20 mM Tris-HCl pH 8, 150 mM NaCl, 2 mM TCEP). Selected peak fractions were ran on an SDS-PAGE gel. Fractions containing both σ^54^ and RNAP core subunits were pooled and concentrated using a 100 kDa MWCO centrifugal filter (Merck Millipore^TM^), supplemented with 10% glycerol and aliquoted into 20 μl volumes, flash frozen in liquid nitrogen and stored at -80 °C.

### Complex formation assays

#### Nucleic acid scaffold preparation

DNA in all experiments used 63 bp DNA from positions -35 to +28. Single stranded DNA was resuspended in 20 mM Tris-HCl buffer and annealed in equimolar amounts in 1× annealing buffer. The mixture was then heated to 95 °C for 5 minutes, and then cooled to 4 °C in 2 °C steps, with 1 minute/step, before being stored at -20 °C. Final concentrations of dsDNA was 100 μM. For DNA-RNA scaffolds, all DNA and RNA oligos were mixed simultaneously, heated and then cooled down as above, and the final concentration was 66.7 μM.

#### Native-PAGE

All native-PAGE assays were run using 4.5% TBE gels in 1× TBE buffer (Invitrogen) running buffer. The recipe for native-PAGE gels and binding buffer used as the loading dye is shown below. Gels were run at 100 V, 4 °C for 100 minutes.

Gels were stained for DNA with SYBR Safe (Thermofisher Scientific™), and protein with InstantBlue (Expedeon^TM^). Final reaction volumes were 20 μl.

To form RNAP-σ^54^-DNA complexes, 250 nM RNAP was mixed with 350 nM DNA for 1 hour at 4 °C in STA buffer (25 mM Tris-Acetate, 8 mM Mg-Acetate, 10 mM KCl, 1 mM TCEP), native-PAGE binding buffer (25% v/v glycerol, 750 mM KCl, 100 mM Tris-HCl pH 8.0, 10 mM MgCl_2_) for 4°C for 1 hour. To form RNAP-σ^54^-DNA-PspF_1-275_ complexes, ADP.AlFx was formed to trap PspF_1-275_ in a hexameric state. The RPc complex was mixed with 1.75 μM PspF_1-275_, 8 mM ADP and 10 mM NaF and incubated for 5 minutes at 37 °C. Following the incubation, the mixture was transferred to ice, and 2 mM AlCl_3_ was added, and then the mixture was incubated for another 12 minutes at 37 °C.

| Gel type | Components (making 5 gels) |
| --- | --- |
| Native-PAGE (4.5% TBE gel) | 7.5 ml 30% Acrylamide (Sigma-Aldrich)<br>5 ml 10× TBE<br>37 ml H<sub>2</sub>O (to 50 ml) |
|  | 0.5 ml 10% ammonium persulfate<br>50 µl Tetramethylethylenediamine |
| Native-PAGE<br>Binding buffer (5×) | 25%v/v Glycerol 750mMKCl 100mM Tris-HCl pH8<br>10 mM MgCl<sub>2</sub> |

### Complex formation for cryoEM studies

#### RPi(-11/-8) complexes

The molar ratios used in gel filtration were identical to the molar ratios used in native-PAGE. 20 μM RNAP-σ^54^ was mixed with 28 μM DNA in Buffer EM1 (20 mM Tris-HCl, 150 mM NaCl, 10 mM MgCl_2_, 1 mM TCEP) for 1 hour at 4 °C, followed by addition of 140 μM of PspF_1-275_, 8 mM ADP and 10 mM NaF. The mixture was incubated at 37 °C for 5 minutes, followed by addition of 2 mM AlCl_3_, and another incubation at 37 °C for 12 minutes. The final volume was 250 μl. The mixture was then centrifuged at 8000 *×g* for 5 minutes at 4 °C before loading onto a Superose 6 10/300 Increase column (Cytiva) equilibrated with Buffer EM1. Selected fractions were analysed by SDS-PAGE to confirm the presence of all protein components, and the fraction at OD_280_ = 200 mAU (∼0.8 μM) on the left shoulder of the peak was used immediately for grid preparation.

#### RPi(-10/-1) complex

RPi(-10/-1) complexes were prepared with identical molar ratios to the above RPi(-11/-8) complexes, except using a final volume of 80 μl and without the gel filtration step. 11.5 μM RNAP-σ^54^ was used, with the same ADP, NaF and AlCl_3_ concentrations as before. Following formation of ADP.AlFx, samples were buffer exchanged into Buffer EM2 (20 mM Tris-HCl, 150 mM KCl, 5 mM MgCl_2_, 5 mM TCEP) using a 0.5 ml Zeba™ 7K MWCO desalting column as per the manufacturers’ protocol. 8 mM CHAPSO was then added immediately before grid preparation.

### Cryo-EM sample preparation

All grids were prepared using a Vitrobot™ Mark IV (FEI) at 4 °C and 100 % humidity, using Grade 595 Vitrobot™ filter paper (Electron Microscopy Sciences). All grids were plunge frozen using liquid ethane and stored in liquid nitrogen.

#### Sample preparation of RPi(-11/-8) complexes

4 μl of the ∼0.8 μM RNAP-σ^54^-DNA-PspF_1-275_ complexes directly from gel filtration were deposited onto 300 mesh holey copper Quantifoil R2/2 grids, that were argon cleaned for 45 seconds (Lambda photometrics^TM^). The blotting parameters were as follows: wait time 30 seconds, blot time 2 seconds and blot force -6.

#### Sample preparation of RPi(-10/-1) complexes

4 μl of complex was deposited onto 300 mesh holey gold C-flat R1.2/1.3 grids, that were plasma cleaned in air for 30 seconds (Harrick Plasma^TM^). The blotting parameters were as follows: wait time 30 seconds, blot time 2 seconds and blot force -8.

#### Data collection

All datasets were collected on a Titan Krios (ThermoFisher Scientific^TM^), operated at 300 kV, equipped with a K3 direct electron detector and a Bioquantam energy filter (Gatan^TM^). Movies were collected at a nominal magnification of 81,000× and a slit width of 20 eV. Data collection was carried out using EPU software (ThermoFisher Scientific^TM^).

| **Sample** | **Pixel Size (Å/pixel)** | **Total Dose (e<sup>-</sup>/Å<sup>2</sup>)** | **Defocus Range (µm)** | **Total Movies** |
| --- | --- | --- | --- | --- |
| RPi(-11/-8) | 1.1 | 50 | -3 to -1 | 11,634 |
| RPi(-10/-1) | 1.1 | 40 | -3 to -1 | 15,534 |

#### Image processing

All image processing steps were carried out in RELION 4.0 (26). Contrast transfer function estimation was carried out with CTFFIND 4 (27). Particle picking in the RPi(-11/-8) dataset was carried out using Gautomatch (https://github.com/JackZhang-Lab/Gautmatch) using RPi(-12/-11) reference projections (28), whereas particles of RPi(-10/-1) dataset were picked with Topaz (29). Detailed image processing pipelines for each dataset are shown in supplementary figures.

## Supplementary Information

**Table S1.** Data collection, image processing, model refinement and statistics.

| Sample |  | Pixel Size<br>(Å/pixel) |  | Total Dose<br>(e<sup>-</sup>/Å<sup>2</sup>) |  | Defocus<br>Range<br>(µm) |  | Total Movies |
| --- | --- | --- | --- | --- | --- | --- | --- | --- |
| RP(-11/-8) |  | 1.1 |  | 50 |  | -3 to -1 |  | 11 634 |
| RP(-10/-1) |  | 1.1 |  | 40 |  | -3 to -1 |  | 15 534 |
| Sample | RP(-11/-8)<br>conformation 1 | RP(-11/-8)<br>conformation 2 | RP(-11/-8)<br>conformation 3 | RP(-11/-8)<br>conformation 4 | RP(-11/-8)<br>conformation 5 | RP(-11/-8)<br>conformation 6 | RP(-<br>10/-1) | pre-RPo<br>state |
| Non-hydrogen<br>atoms | 26960 | 26819 | 26953 | 26928 | 26832 | 26903 | 26931 | 19635 |
| Protein<br>residues | 5154 | 5125 | 5153 | 5148 | 5128 | 5143 | 5165 | 3581 |
| Nucleic acid<br>residues | 68 | 68 | 68 | 68 | 68 | 68 | 64 | 87 |
| Ligands | 5 Mg<sup>2+</sup> | 5 Mg<sup>2+</sup> | 5 Mg<sup>2+</sup> | 5 Mg<sup>2+</sup> | 5 Mg<sup>2+</sup> | 5 Mg<sup>2+</sup> | 5 Mg<sup>2+</sup> | 1 Mg<sup>2+</sup> |
|  | 5 ADP | 5 ADP | 5 ADP | 5 ADP | 5 ADP | 5 ADP | 5 ADP |  |
|  | 5 AF3 | 5 AF3 | 5 AF3 | 5 AF3 | 5 AF3 | 5 AF3 | 5 AF3 |  |
| B factors (Å<sup>2</sup>) |  |  |  |  |  |  |  |  |
| Protein | 191.59 | 273.75 | 225.37 | 397.12 | 595.25 | 882.62 | 349.09 | 105.54 |
| Nucleic acid | 461.16 | 289.8 | 438.31 | 640.15 | 235.94 | 284.5 | 385.72 | 296.67 |
| Ligands | 121.29 | 563.98 | 171.92 | 585.45 | 307.84 | 706.08 | 344.07 | 326.69 |
| R.m.s.<br>deviations |  |  |  |  |  |  |  |  |
| Bond lengths<br>(Å) | 0.007 | 0.002 | 0.003 | 0.01 | 0.002 | 0.011 | 0.002 | 0.004 |
| Bond angles<br>(°) | 1.032 | 0.641 | 0.655 | 0.758 | 0.653 | 0.904 | 0.543 | 0.857 |
| MolProbity<br>score | 2.52 | 1.52 | 1.68 | 2.11 | 2.09 | 2.42 | 1.83 | 2.09 |
| Clashscore | 23.55 | 3.53 | 4.7 | 11.71 | 10.91 | 19.19 | 6.79 | 8.4 |
| Poor rotamers<br>(%) | - | - | - | - | - | - | - | - |
| Ramachandran<br>plot |  |  |  |  |  |  |  |  |
| Favored (%) | 85 | 93.84 | 93.31 | 90.73 | 90.57 | 0.2 | 92.78 | 86.1 |
| Disallowed (%) | 0.43 | 0.04 | 0 | 0.12 | 0.08 | 0.86 | 0.02 | 0.84 |

**Fig. S1.**
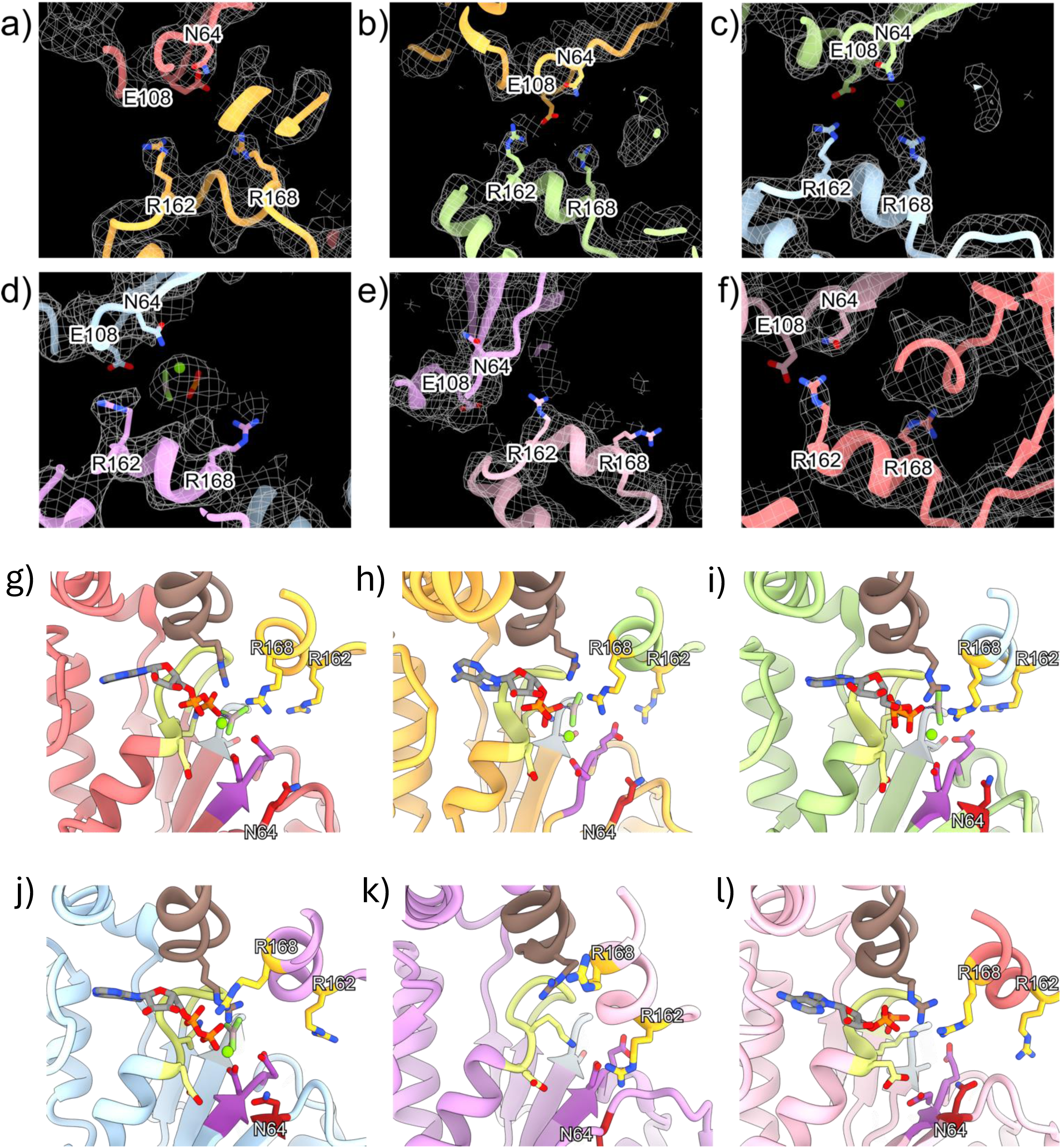
nucleotide binding pockets of each protomer within the hexamer. (a)-(f) electron density for the key catalytic residues (g)-(l) positions of catalytic residues relative to nucleotides.

**Fig. S2.**
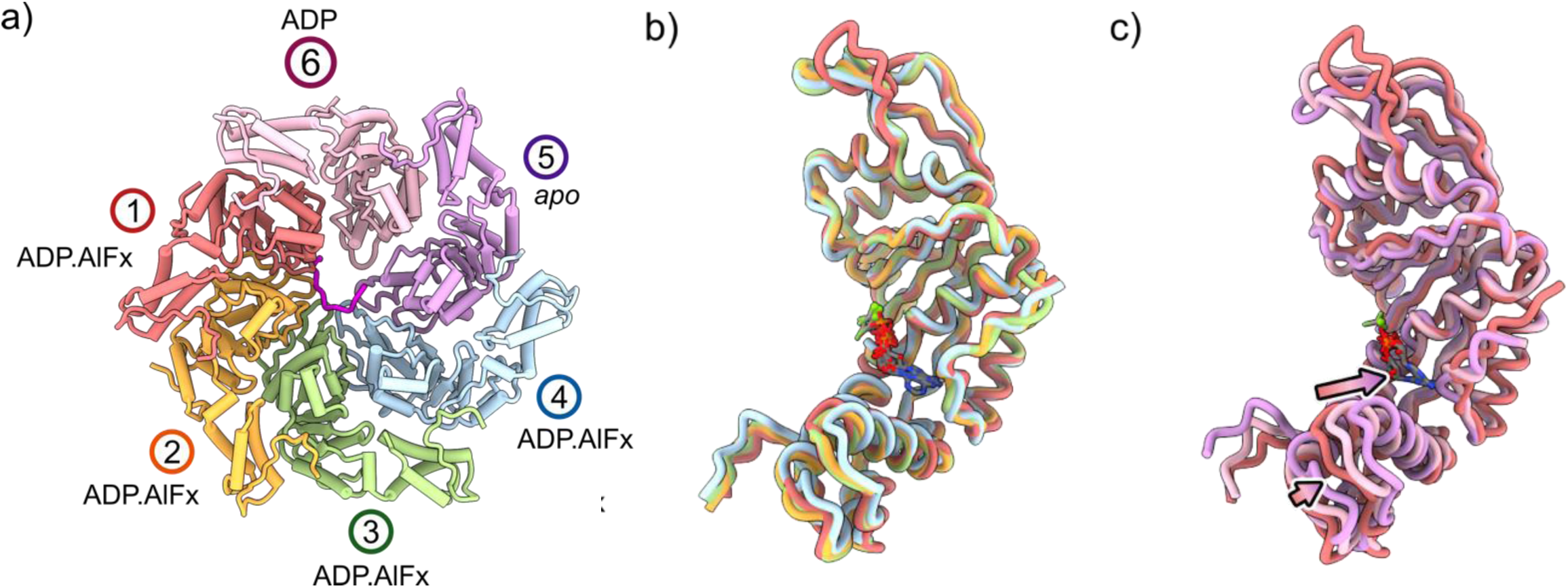
Comparisons of PspF_1-275_ protomers within the RPi(-12/-11) complex. (a) hexamer and their nucleotide states of each protomer. (b) Protomers 1-4 are similar in conformation. (c) comparisons of Protomer 1, protomer 6 and protomer 5 shows the conformational changes from ATP hydrolysis (ATP to ADP) and nucleotide release (ADP to nucleotide free).

**Fig. S3.**
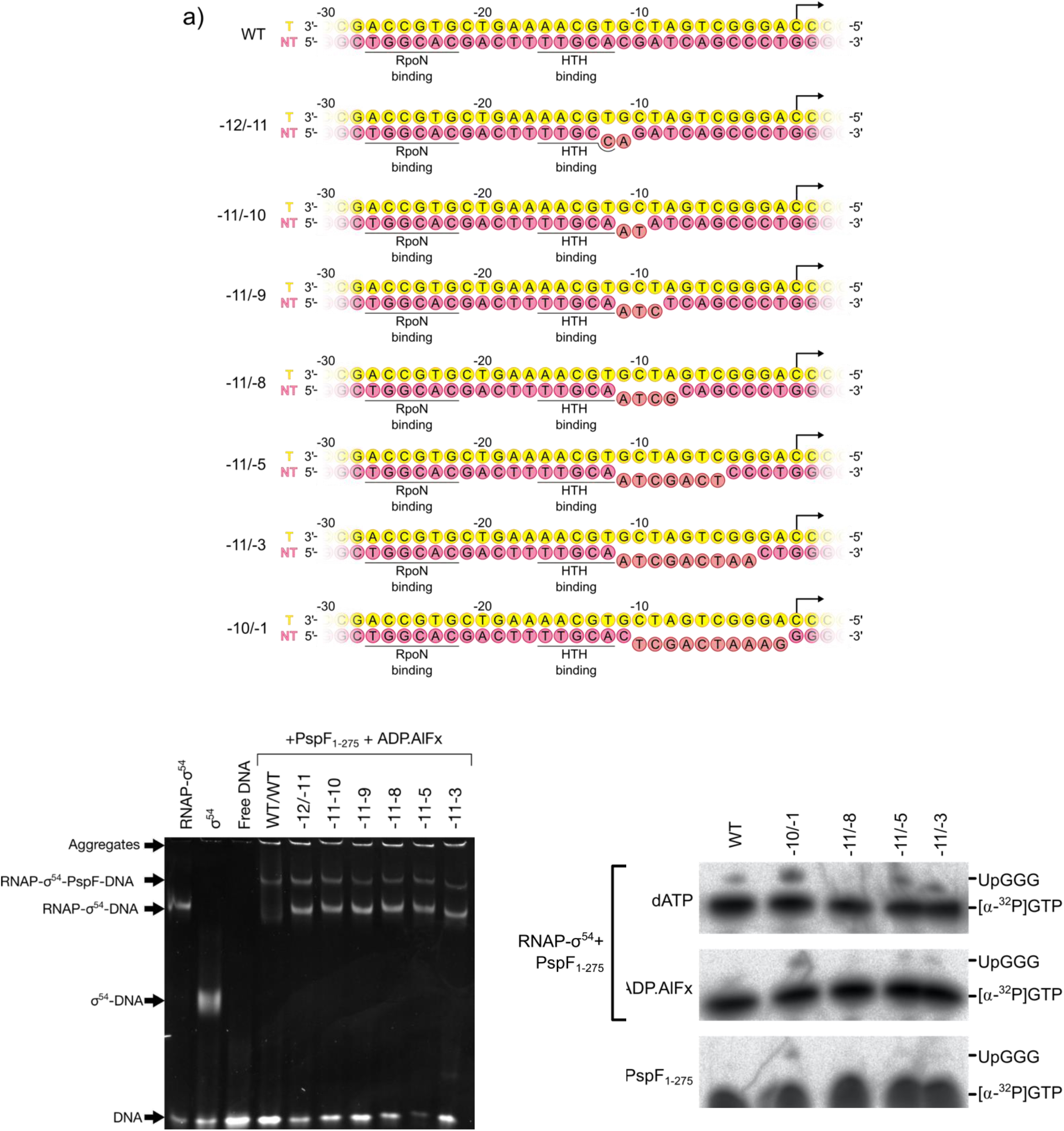
Biochemical characterizations of intermediate complexes. (a) DNA susbtrates used to mimick progressively opening up DNA, (b) Native-PAGE gel of RNAP-σ^54^-PspF_1-275_ complexes trapped with ADP.AlFx and varying degrees of DNA mismatch. Lane labels indicate mismatched bases. For example, -11-9 indicates mismatched bases between -11 and -9. (c) ability of initiating transcription transcription in the presence of ADP.AlFx or dATP.

**Fig. S4.**
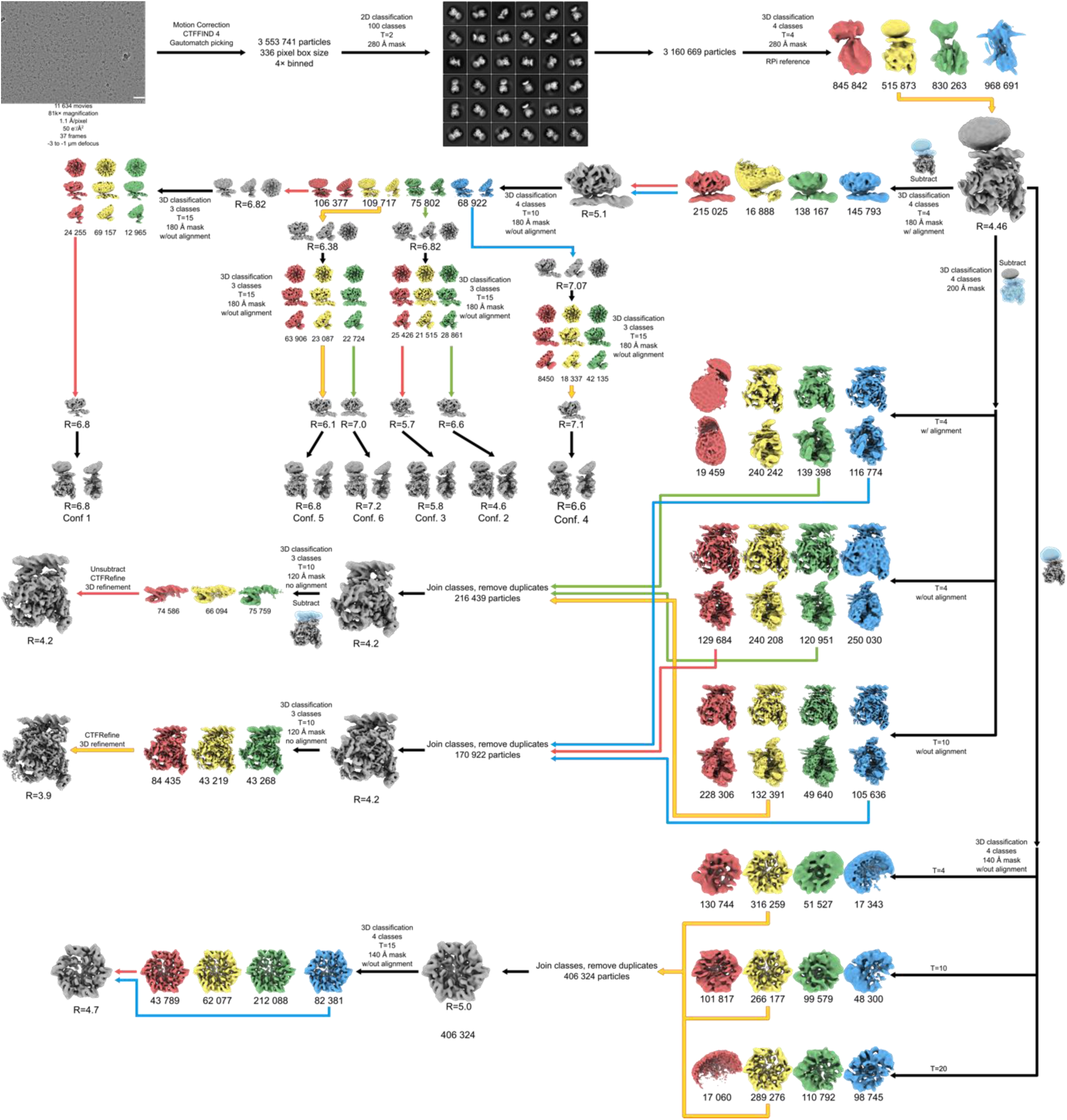
image processing flow chat for RPi (-11/-8) complex.

**Fig. S5.**
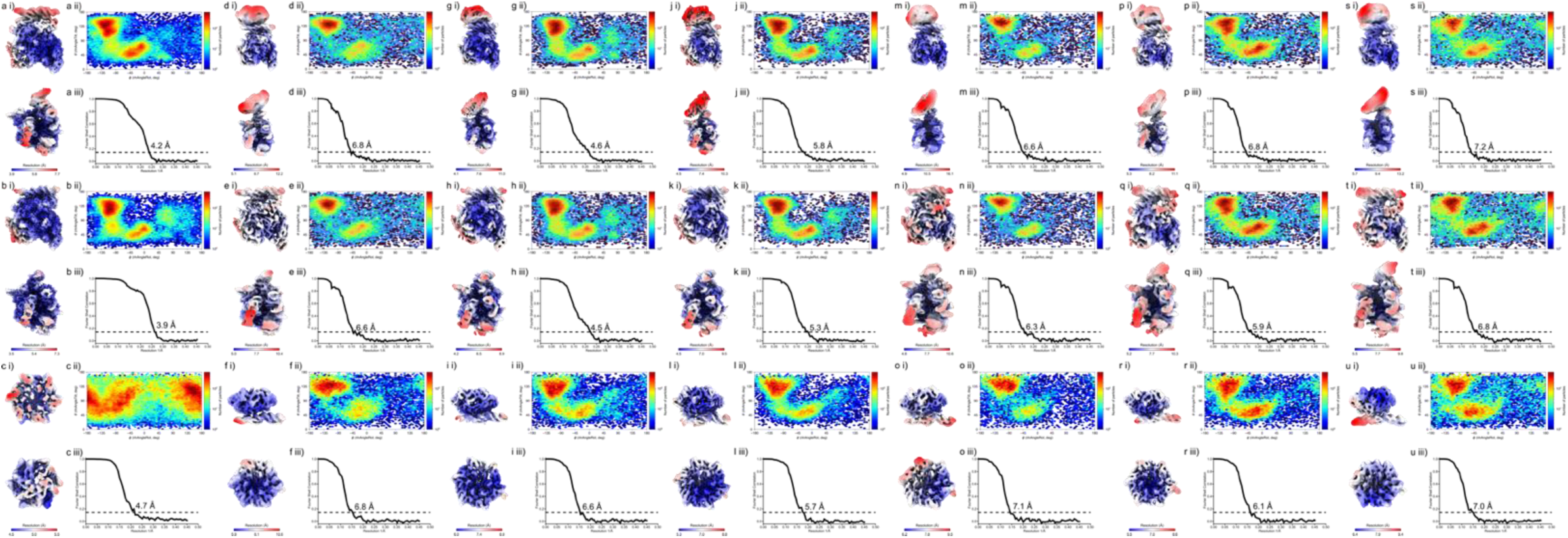
Local resolution map, angular distribution of particles and FSC curves of various reconstructions from RPi(-11/-8) complex. a) open clamp and b) closed clamp c) and PspF maps obtained from subtracting the initial consensus map. Conformations 1 (d-f), 2 (g-i), 3 (j-l), 4 (m-o), 5 (p-r) and 6 (s-u) with the maps on the first row showing local resolution estimation of the global refinement, the second row obtained from subtracting the RNAP-σ^54^-DNA density and running refinement and local resolution estimation, and the 3^rd^ row obtained from subtracting the PspF-σ^54^(excluding region II)-DNA density and running refinement and local resolution estimation. The subtracted refined maps were then used to create the composite maps shown in Supplementary figure 6. For each panel, i) shows the local resolution map ii) shows the angular distribution and iii) shows the FSC curves.

**Fig. S6.**
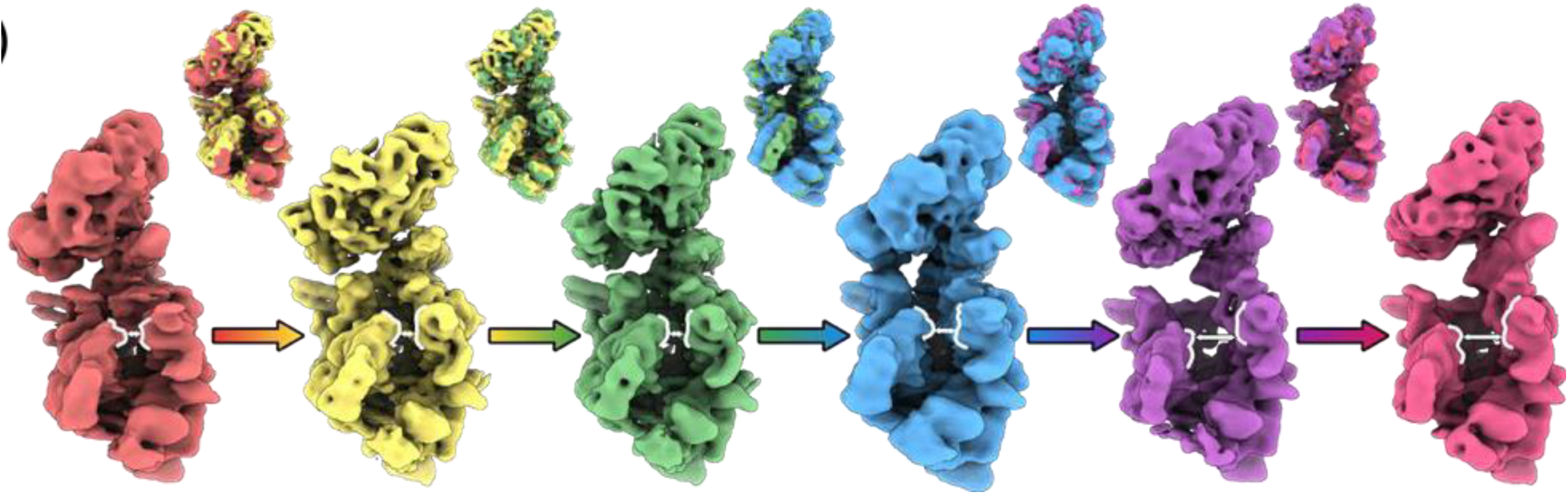
Conformational heterogeneity of RPi (-11/-8), six well resolved conformations showing varying degrees of PspF_1-275_ tilt and clamp opening.

**Fig. S7.**
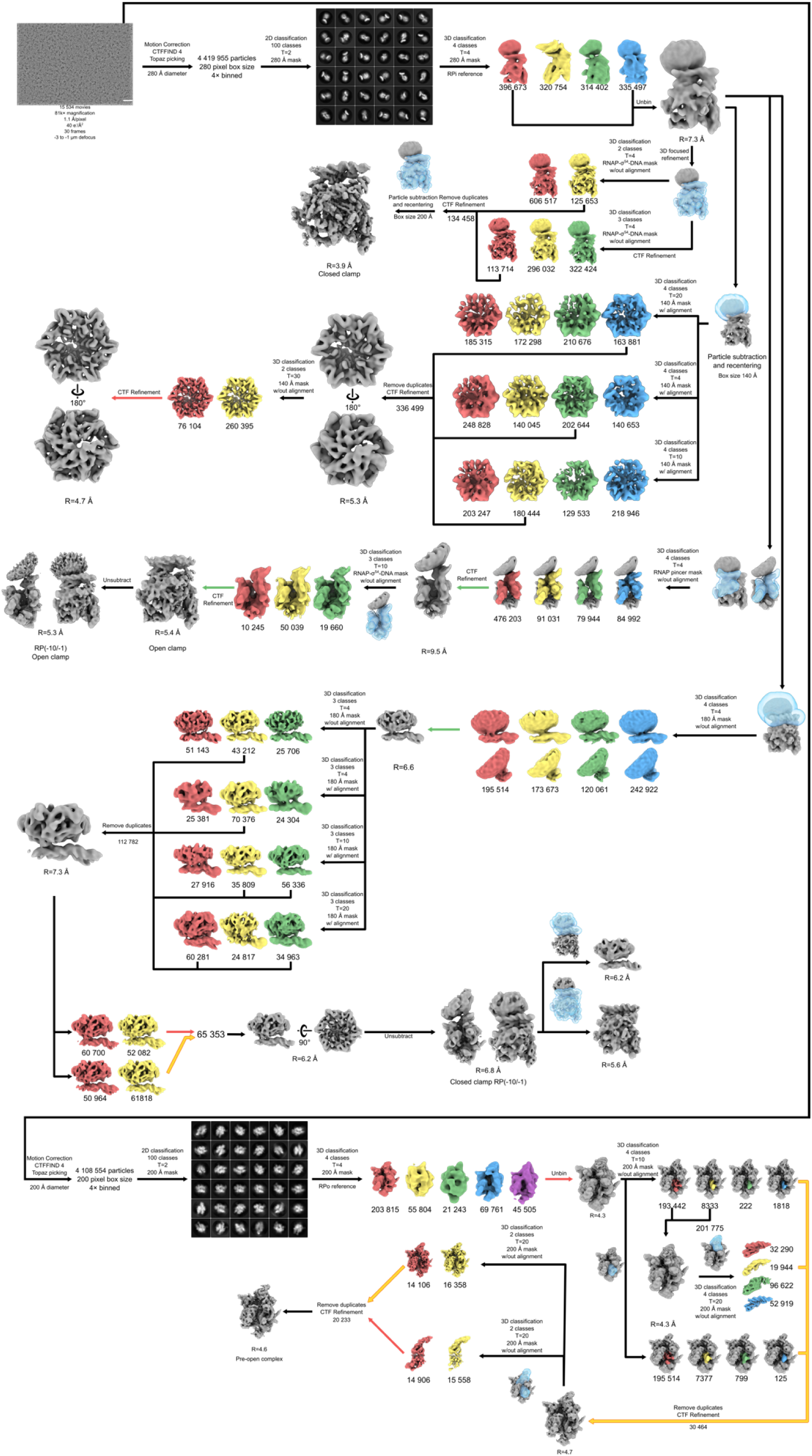
Image processing flow chat of RPi (-10/-1) complex.

**Fig. S8.**
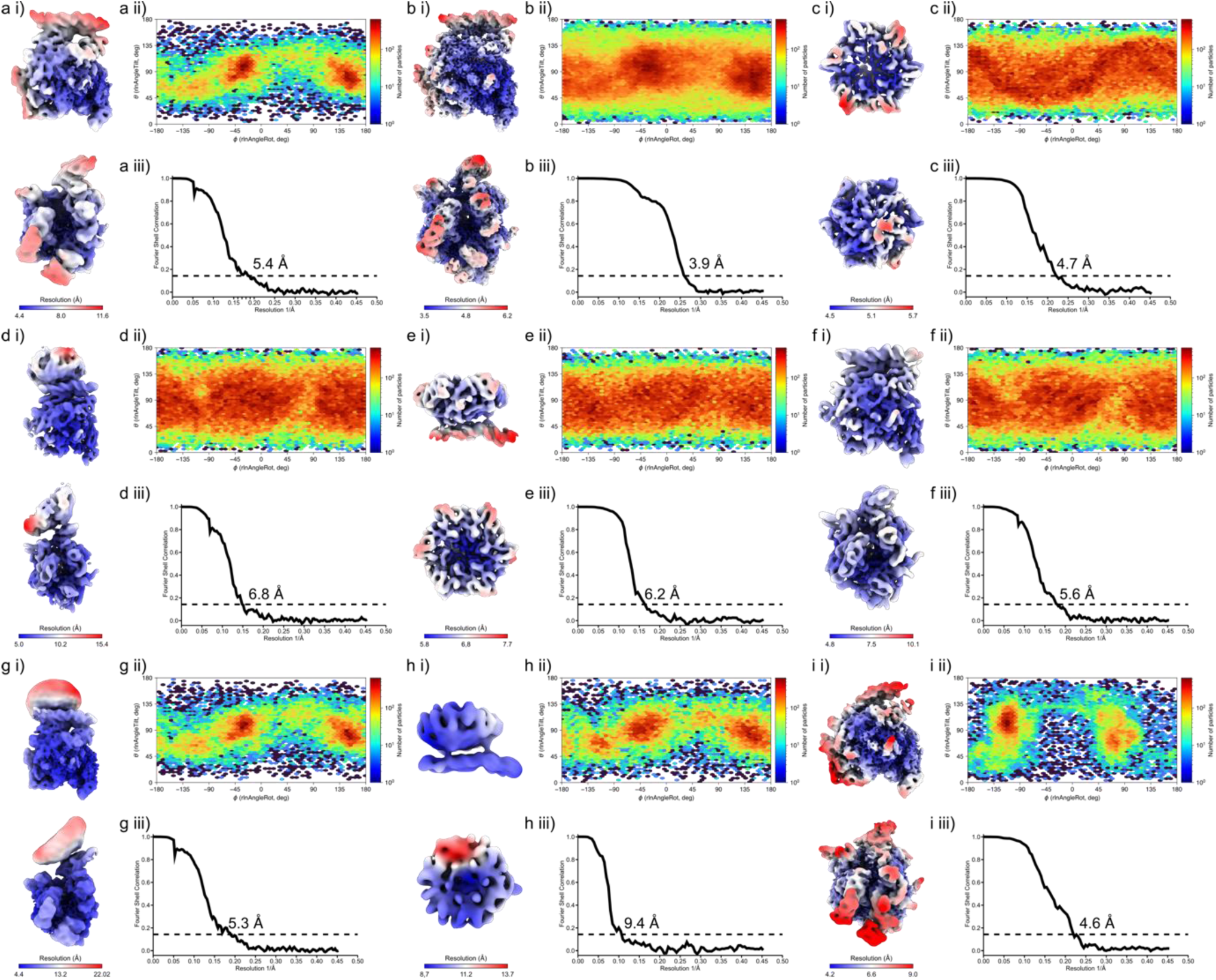
Local resolution map, angular distribution of particles and FSC curves of various reconstructions from RPi(-10/-1) complex. a) open clamp and b) closed clamp c) and PspF maps obtained from subtracting the initial consensus map. d-f) closed clamp conformation obtained from reverting back to the original particle, whilst retaining the angular assignments the PspF-σ^54^(excluding region II)-DNA density classifications and running global refinement and local resolution estimation (d), followed by focused classification, refinement and local resolution estimation of the PspF-σ^54^(excluding region II)-DNA and RNAP-σ^54^-DNA. g) Open clamp conformation obtained unsubtracting the RNAP β’ clamp and β lobe - σ^54^-DNA density classifications and running global refinement and local resolution estimation (g), followed by focused classification, refinement and local resolution estimation followed by focused classification, refinement and local resolution estimation of the PspF-σ^54^(excluding region II) (h). i) Open complex densities. For each panel, i) shows the local resolution map ii) shows the angular distribution and iii) shows the FSC curves.

**Fig. S9.**
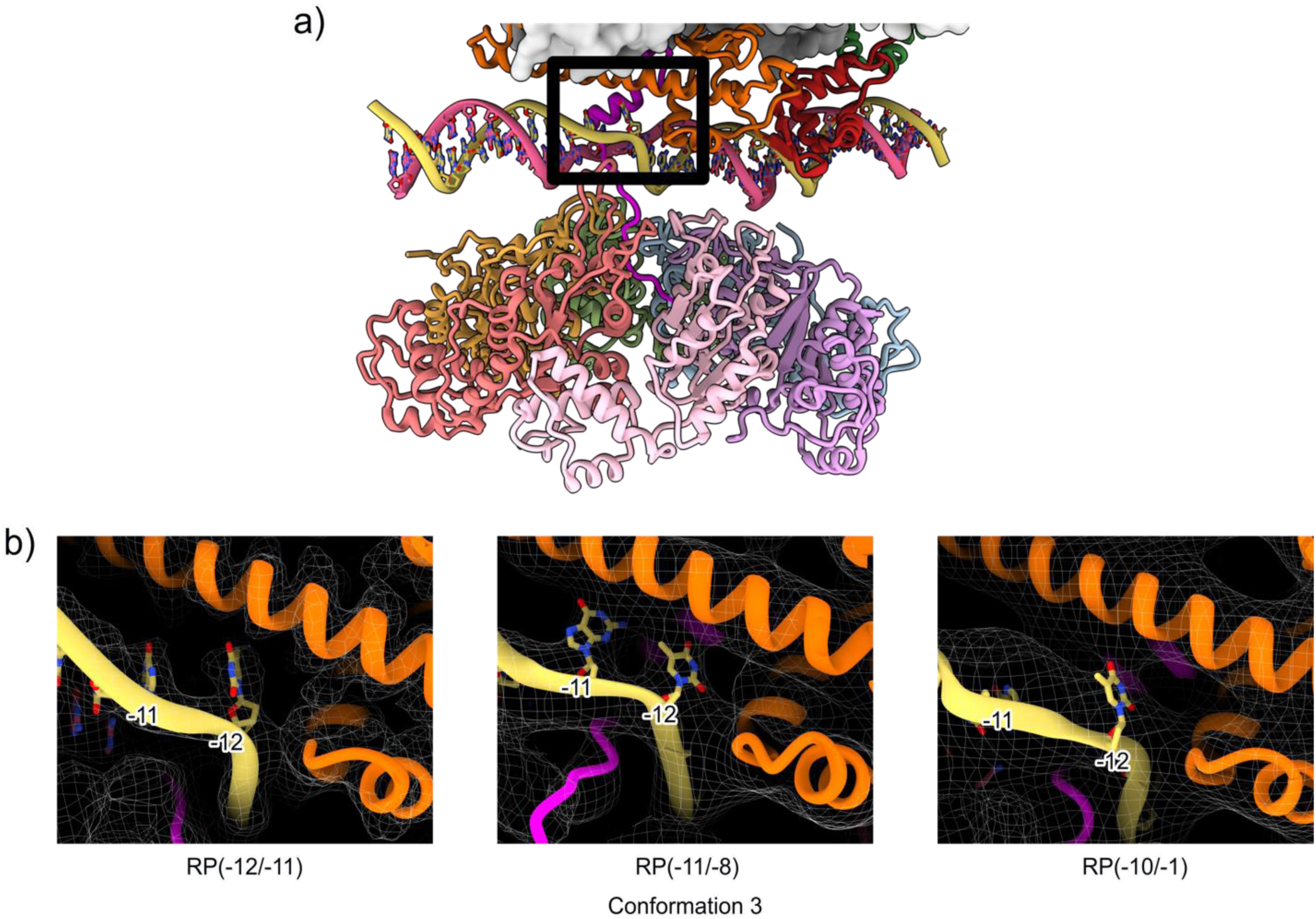
Density of DNA at -12/-11 for RPi (-12/-11), RPi(-11/-8) and RPi(-10/-1).

**Fig. S10.**
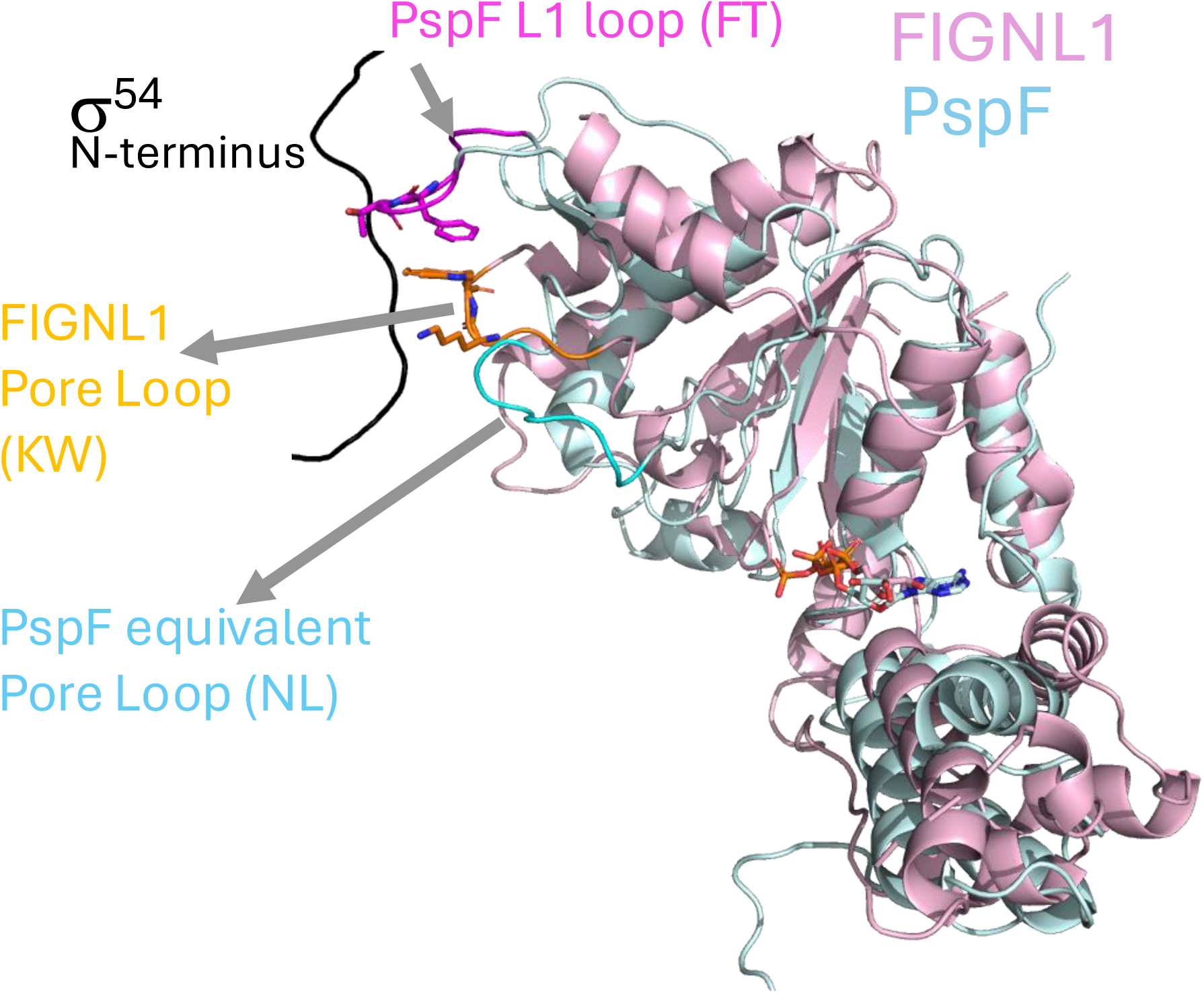
Comparisons of the AAA+ domains of PspF_1-275_ with that of FIGNL1. FIGNL1 pore loops are shown to interact and translocate substrates. PspF does not have residues which are conserved among AAA+ translocases. Instead L1 loops are positioned similarly to pore loops and perform similar roles.

